# mRNA transcription in skeletal muscle drives growth and determines nuclear accretion

**DOI:** 10.64898/2025.12.29.696842

**Authors:** Fabian Montecino-Morales, Cristofer Calvo, Casey O. Swoboda, Alena Bruening, Alyssa A.W. Cramer, Xin Lin, Jennifer L. Martin, Ramzi J. Khairallah, Yarui Diao, Vikram Prasad, Douglas P. Millay

## Abstract

Multinucleation of skeletal muscle cells (myofibers) is a determinant of size and fundamental for function. While it is established that myofibers need to accrue adequate numbers of nuclei for optimal growth, the molecular circuitry linking myonuclei to growth and why myofibers need additional nuclei remains unknown. We found that growth is still possible after restriction of nuclear content in myofibers and this was associated with increased levels of RNA Polymerase II (*Polr2a*) leading to elevated mRNA content. Through development of a genetic mouse model where endogenous *Polr2a* is upregulated in myofibers, we established that increased transcriptional output is sufficient to drive functional growth of myofibers. Notably, we discovered that *Polr2a* overexpression curtails the need for additional nuclei for myofiber growth. These data reveal a previously neglected driver of functional muscle growth and highlight that increasing *Polr2a*-mediated transcription from the vast numbers of nuclei within myofibers could be leveraged to combat muscle wasting conditions.

## Introduction

Adult skeletal muscle cells (myofibers) are amongst the largest cells in mammals with volumes multiple log orders larger than other large cells. Myofibers are multinucleated, terminally differentiated syncytial cells that accumulate vast numbers of nuclei through cell fusion of muscle progenitor cells (*1, 2*). The most pronounced increase in myofiber sizes occurs during postnatal and pubertal developmental phases, but it is noteworthy that growth of myofibers occurs after the majority of nuclear accretion has occurred (*2–4*). In response to external stimuli, adult myofibers can accrue additional nuclei through fusion with activated satellite cells (muscle stem cells) prior to any changes in myofiber size (*5*), and this nuclear accretion, in adequate numbers, is critical for sustained functional growth (*1, 6–8*). The robust growth of myofibers that follows accrual of nuclei suggests that the biosynthetic output from these resident myonuclei facilitates the observed growth. It is reasonable to infer that this increase in myonuclear output should encompass an increase in mRNA abundance. However, direct experimental evidence for such augmented mRNA transcription remains sparse and much of the information on myofiber growth focuses on the expansion of translational capacity (*9–16*).

Evidence that mRNA content within myofibers may be attuned to growth comes from a mouse model where myonuclear accretion was restricted during development (*17*). In this model, myofibers partially mitigate the negative effects of reduced DNA content on growth through an increase in mRNA content. This finding is consistent with observations that mRNA content scales with cell size across species and various cell types (*18–22*) and where the concentration of biosynthetic products, including mRNA, remains largely stable as cells grow and volumes enlarge in mononucleated, proliferative cells (*23–26*). Mechanistically, mRNA scaling in mononuclear cells involves increased transcription resulting from elevated expression, and genome loading of the multi-subunit RNA Polymerase II (RNAPII) enzyme complex (*27, 28*). Despite these advances in our understanding of mRNA scaling with cell size, a key question that has remained unanswered is that of causation, whether increasing mRNA transcriptional capacity of nuclei, and mRNA content within cells are sufficient to drive increases in cell size.

Here, we directly test the links between myonuclear mRNA transcriptional output and myofiber growth. We demonstrate that while multinucleation serves a critical function in establishment of myofiber size, enhancing their biosynthetic output through upregulation of the *Polr2a* gene (Rpb1 protein), a key subunit of RNAPII, can be exploited for growth. Additionally, we found that activation of RNAPII in myofibers reduces the need for growth-associated nuclear accretion. This unexpected observation suggests the existence of a feedback mechanism where regulation of mRNA transcriptional output in myofibers impacts muscle progenitors and accretion. Since current efforts to optimize muscle growth are focused on direct regulation of proteostasis and activation of satellite cells to fuse and contribute additional DNA content to myofibers, we anticipate the paradigm revealed here will provide additional approaches to regulate myofiber sizes, which is critical for the health and longevity of mammals (*29–32*). Overall, we establish that the transcriptional activity of resident myonuclei can drive muscle growth and identify a fundamental cell biological principle that underlies the need for myofibers to accrue nuclei.

## Results

### Enhanced transcriptional dynamics through elevated RNA polymerase II in myofibers with fewer myonuclei

The mouse model where myonuclear accretion was abrogated during development (Δ2w mice) was achieved through genetic deletion of Myomaker in muscle progenitors (*17*), resulting in myofibers with 50% fewer nuclei. Our previous analyses showed that Δ2w mice exhibit overall smaller myofibers but volumes per myonucleus are increased compared to myofibers with their full complement of nuclei. Based on analysis of whole muscle, our results suggested that mRNA content per myonucleus was increased. We sought to understand mRNA content and transcription at the myofiber and myonuclear levels in Δ2w mice to identify specific mechanisms for regulation of myofiber volumes. Through smRNA-FISH with a probe for polyA mRNA, we observed that myofibers from Δ2w mice exhibited similar amounts of total mRNA but since there are fewer nuclei there is an increase in mRNA concentration (total mRNA normalized to number of nuclei) (Fig. 1A). These data confirm the finding that myofibers with fewer myonuclei have elevated mRNA per nucleus. To test whether multinucleation itself can impact mRNA concentration, we generated artificial syncytiums by forcing fibroblasts to fuse through the expression of the muscle fusogens Myomaker and Myomerger (*33–35*) (fig. S1A). We found that with increasing amounts of multinucleation there was reduced polyA mRNA per nucleus (fig. S1B). These data suggest that in multinucleated cells, transcriptional output of individual nuclei falls as nuclear numbers increase. This means that transcriptional capacity of nuclei within syncytia, such as myofibers, might not be utilized to the extent it is in mononucleated cells; such underutilization raises the possibility that some transcriptional capacity remains available for mobilization in nuclei within myofibers.

**Figure 1.**
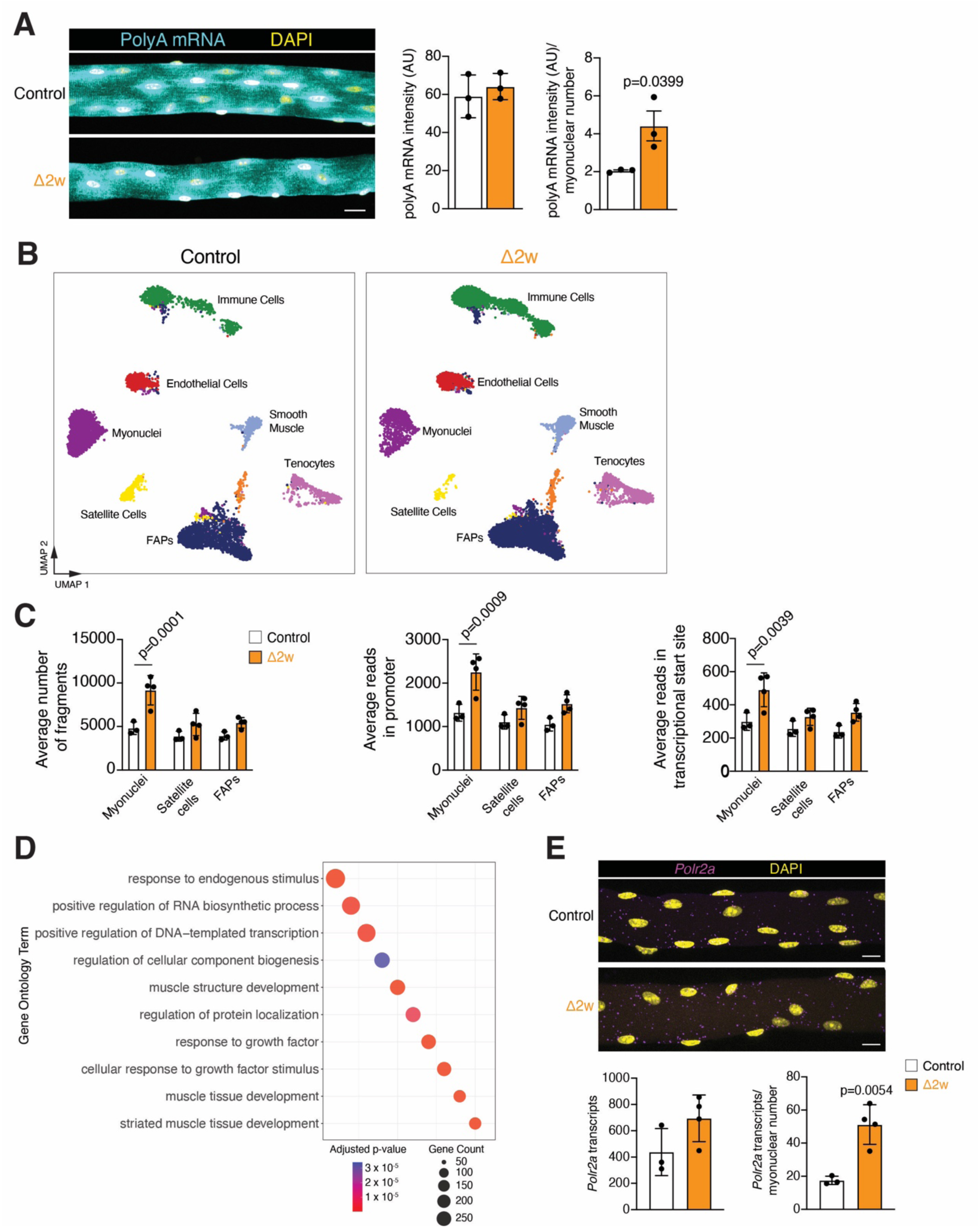
Myofibers with fewer myonuclei exhibit characteristics of increased transcription. (A) Representative images from polyA mRNA smRNA-FISH on control and Δ2w EDL myofibers at postnatal day 42. Scale bar: 10 μm. Quantification of polyA mRNA intensity and normalized by the number of myonuclei (n=3). (B) UMAP from snATAC-sequencing of control and Δ2w muscle at postnatal day 42. (C) Average number of total fragments (left), average reads in promoter (center), average reads in transcriptional start site (right) in myonuclei, satellite cells, and fibroadipocyte progenitors (FAPs) from control and Δ2w mice (n=3-4). (D) Gene ontology enrichment analysis of genes nearest to differentially accessible peaks increased in Δ2w mice from snATAC-sequencing. (E) Representative images for *Polr2a* smRNA-FISH on isolated myofibers from control and Δ2w mice at postnatal day 42. Scale bar: 10 μm. Quantification of transcript number for *Polr2a* and normalized by nuclei number (n=3-4). Data are presented as mean ± SD. Statistical tests used were (A), (E) unpaired t-test; (C) two-way ANOVA with Šídák’s correction for multiple comparisons.

To identify possible mechanisms leading to increased mRNA concentration in Δ2w muscle, we directly analyzed the rate of transcription. Nascent RNA was labelled by administering 5-Ethynyl Uridine (EU) to control or Δ2w mice, followed by a chase of 6 hours, at which point muscle was harvested, total RNA isolated, and EU-labelled RNA was pulled down. From the total RNA pool, quantitative RT-PCR for muscle-specific sarcomeric genes *Myh4* and *Acta1* followed by normalization to myonuclear number revealed an increased expression of these genes in Δ2w samples (Fig. S1C). Increases in the EU-labelled RNA pool was also detected revealing an increased rate of transcription in Δ2w muscle (Fig. S1C).

We then utilized snATAC-seq to determine if increased transcription was associated with alterations in chromatin accessibility. Unbiased clustering revealed all major cell populations from control and Δ2w skeletal muscle (Fig. 1B) and these cell populations were identified using gene accessibility scores for known marker genes (fig. S2). Despite the expected reduced number of myonuclei in the Δ2w sample, we observed an increased average number of DNA fragments from myonuclei, which is an indicator of more accessible chromatin (Fig. 1C). This increased number of fragments included reads in promoter and transcriptional start site regions and is only observed in Δ2w myonuclei but not satellite cells or fibroadipocyte progenitors (FAPs) (Fig. 1C). These data indicate that the increased transcription observed in Δ2w myonuclei was associated with more open and accessible chromatin.

We performed gene ontology analysis on identified genes nearest to the differentially accessible peaks from the snATAC-seq data. Gene ontology of the increased genes found in Δ2w myonuclei revealed categories of positive regulation of RNA biosynthesis, DNA transcription, and muscle development (Fig. 1D), overall consistent with enhanced transcriptional output. We analyzed the expression of the *Polr2a* gene that encodes for Rpb1, the major subunit of the RNA polymerase II complex (*36*). smRNA-FISH on isolated myofibers for *Polr2a* mRNA levels was not significantly altered despite Δ2w myofiber volumes being smaller, indicating an increase in overall abundance (Fig. 1E). There was a significant increase in concentration (normalized by myonuclear number) of *Polr2a* mRNA in Δ2w mice (Fig. 1E). These results suggest that the RNAPII complex is upregulated in Δ2w myofibers. This provides one explanation for how these myofibers, despite having fewer resident myonuclei, alleviate the effects of lower DNA content on size by increasing mRNA transcription and content. Based on these findings, and evidence from artificial syncytia, we hypothesized that *Polr2a* levels and mRNA transcriptional output from resident myonuclei could be a tunable component to drive muscle growth.

### Development of a system to increase myonuclear transcription *in vivo*

We sought to develop a mouse model with increased transcriptional output of nuclei within myofibers. We utilized a target gene activation (TGA) dead-guide RNA-Cas9 (dgRNA-Cas9) system (*37*) to upregulate the expression of endogenous *Polr2a* in mouse skeletal muscle. In this system, the dgRNA is shorter (15 base pairs) than normal and also contains an MS2 binding site. Cas9 and the short dgRNA bind to the transcriptional start site of a target gene but this does not result in cutting of the DNA. The system also contains the transcriptional activation complex MS2:P65:HSF1 (MPH), where the MS2 subunit binds to the MS2 binding site on the dgRNA bringing the transcriptional activators P65:HSF1 in proximity to the target gene, thus activating gene transcription. We designed a dgRNA predicted to bind upstream of the transcriptional start site of the *Polr2a* gene and packaged this along with the MPH complex (driven by CAG promoter) into MyoAAV (Fig. S3A), which has high tropism for skeletal myofibers (*38*). To test the system, MyoAAV-CAG-MPH-*Polr2a* dgRNA or MyoAAV-GFP (control) was intramuscularly injected into tibialis anterior muscles of *Rosa26*^Cas9^ mice. We observed an increase in *Polr2a* transcripts and polyA mRNA levels (Fig. S3B) indicating increased transcription in myofibers. MyoAAV-CAG-MPH-*Polr2a* dgRNA also upregulated *Polr2a* compared to a MyoAAV-CAG-MPH-scrambled dgRNA (Fig. S3C), which still expresses the transcriptional coactivator MPH complex indicating that the MPH complex alone is not sufficient for gene activation. Because expression of Cas9 in the *Rosa26*^Cas9^ mouse is ubiquitous and the constructs utilize the CAG promoter and MyoAAV can transduce around 10-20% of satellite cells (*38, 39*), we could also be activating *Polr2a* in non-myofibers. Thus, we generated additional constructs where MPH is driven by the myofiber-specific creatine kinase (CK8e) promoter (*40*) (Fig. S3D). Delivery of MyoAAV-CK8e-MPH-*Polr2a* dgRNA also upregulated *Polr2a* when compared with MyoAAV-CK8e-MPH-scrambled dgRNA (Fig. S3D), and this level of increase was comparable to the upregulation observed with MyoAAV-CAG-MPH-*Polr2a* dgRNA. In addition, upon upregulation of *Polr2a* in this muscle-specific system, we detected increased levels of its protein product, Rpb1 (Fig. S3E). Overall, these data demonstrate the development of a system that allows us to interrogate the effects of increased mRNA transcription, through the upregulation of *Polr2a*, in skeletal myofibers.

We next characterized the nature of increased transcription in myofibers upon upregulation of *Polr2a*. We chose to analyze abundance of transcripts through smRNA-FISH on muscle sections and focused on sarcomeric genes, a major component of myofiber transcriptomes. We performed smRNA-FISH on muscle sections after delivery of MyoAAV-CK8e-MPH-*Polr2a* dgRNA or MyoAAV-CK8e-MPH-scrambled dgRNA (Fig. 2A). *Myh4*, *Myh1*, and *Ttn* all exhibited increased abundance in myofibers that received the *Polr2a* dgRNA (Fig. 2B). We also detected increased signal for the polyA mRNA probe, but there was no increase in the ribosomal RNA 47S pre-rRNA (Fig. 2C) indicating that the increased transcription is biased towards mRNA as expected given the function of RNAPII. These data suggest that *Polr2a* activation in myofibers augments transcription of genes, such as sarcomeric genes, that are normally expressed in muscle.

**Figure 2.**
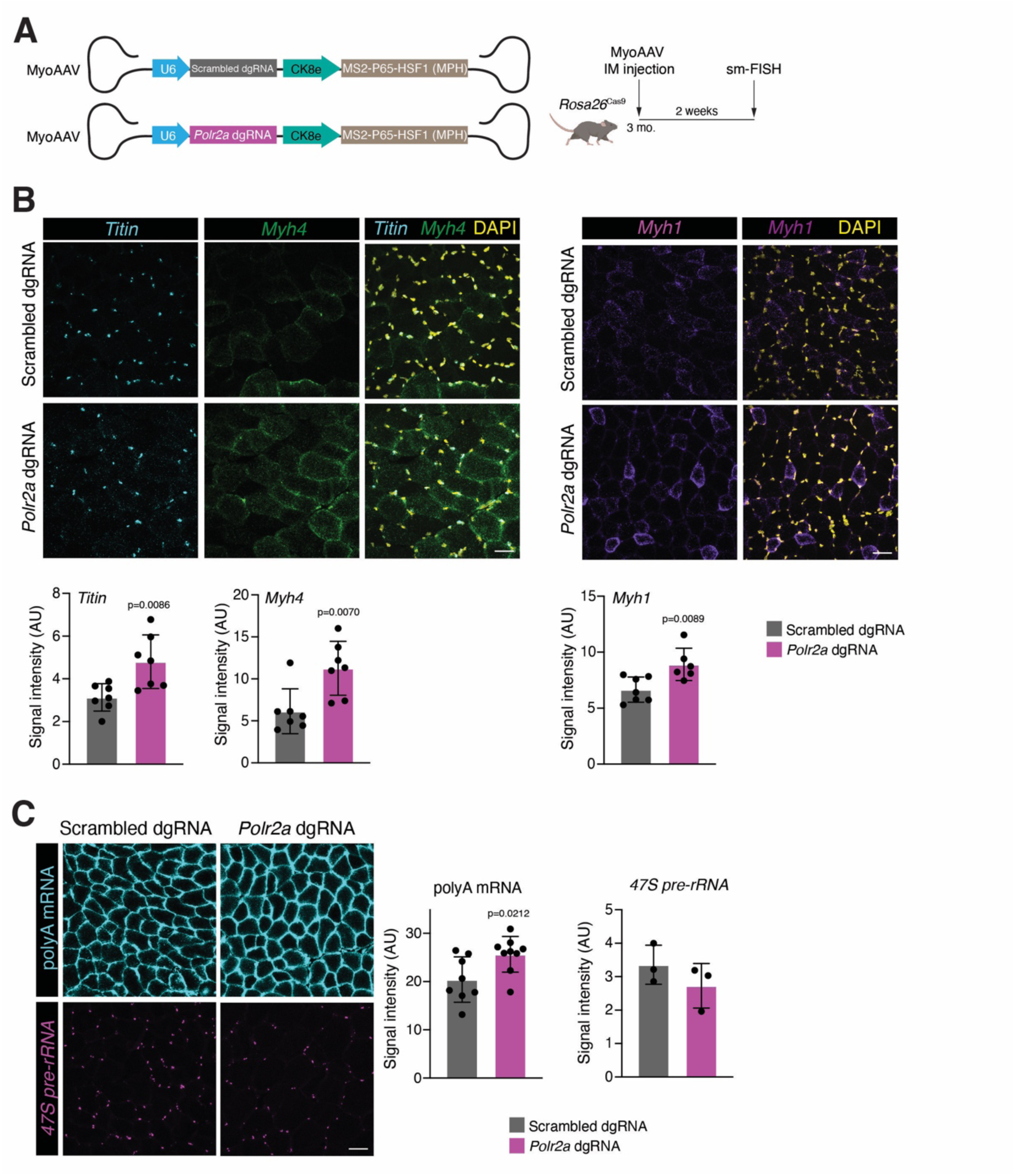
*Polr2a* upregulation increases RNA polymerase II-dependent transcription in myofibers. (A) The indicated MyoAAVs were intramuscularly delivered to the tibialis anterior (TA) of *Rosa26*^Cas9^ mice. TAs were analyzed two weeks after transduction. (B) Representative smRNA-FISH images for *Myh4, Ttn,* and *Myh1* on TA cryosections from adult *Rosa26*^Cas9^ mice transduced with scrambled or *Polr2a* dgRNA. Quantification of signal intensity is shown on the bottom (n=7). Scale bar: 20 μm. (C) Representative smRNA-FISH images for polyA mRNA and 47S pre-rRNA. Quantification of signal intensity quantification is shown on the right (polyA mRNA n=8-9; 47S pre-rRNA n=3). Scale bar: 20 μm. Data are presented as mean ± SD. Statistical tests used were (B) *Myh1*, *Titin*, (C) unpaired t-test; (B) *Myh4* Mann-Whitney test.

### Elevated myonuclear transcription drives functional muscle growth

Having established a system to increase RNAPII-dependent transcription and mRNA levels in myofibers, we wanted to test if elevation of transcription caused an increase in size. After delivery of MyoAAV-CK8e-MPH-*Polr2a* dgRNA or MyoAAV-CK8e-MPH-scrambled dgRNA, we observed increased cross-sectional area of myofibers in muscles that had elevated *Polr2a* (Fig. 3A). Of note, there is efficient transduction of myofibers since we observed increased *Polr2a* in the majority of myofibers (Fig. 3B). In an independent experiment, we also observed myofiber growth in adult muscles transduced with MyoAAV-CAG-MPH-*Polr2a* dgRNA compared to control vectors MyoAAV-CMV-GFP or MyoAAV-CAG-MPH-scrambled dgRNA (fig. S4A). The growth of adult EDL myofibers, which are transduced by intramuscular injection of tibialis anterior muscles, after *Polr2a* activation was not associated with an increase in myonuclear numbers (fig. S4B). However, we observed increased cell size in these isolated myofibers (fig. S4B), indicating that growth associated with *Polr2a* upregulation does not induce nuclear accretion. To confirm the absence of an increase in myonuclear numbers, we administered EdU which will label satellite cells if they are activated to proliferate, prior to differentiation and fusion. This label can then be tracked into myofibers as an indicator of fusion (*41*). We detected minimal EdU^+^ myonuclei in both MyoAAV-CMV-GFP or MyoAAV-CAG-MPH-*Polr2a* dgRNA transduced myofibers (fig. S4C). These data indicate that the *Polr2a* gene activation system does not precociously induce fusion of activated satellite cells, and that the increase in myofiber sizes is independent of an increase in DNA content through fusion with muscle progenitor cells. Thus, these data strongly suggest that an increase in myonuclear transcription achieved by *Polr2a* overexpression sufficiently augments biosynthetic output of resident nuclei within myofibers to expand cytoplasmic volumes.

**Figure 3.**
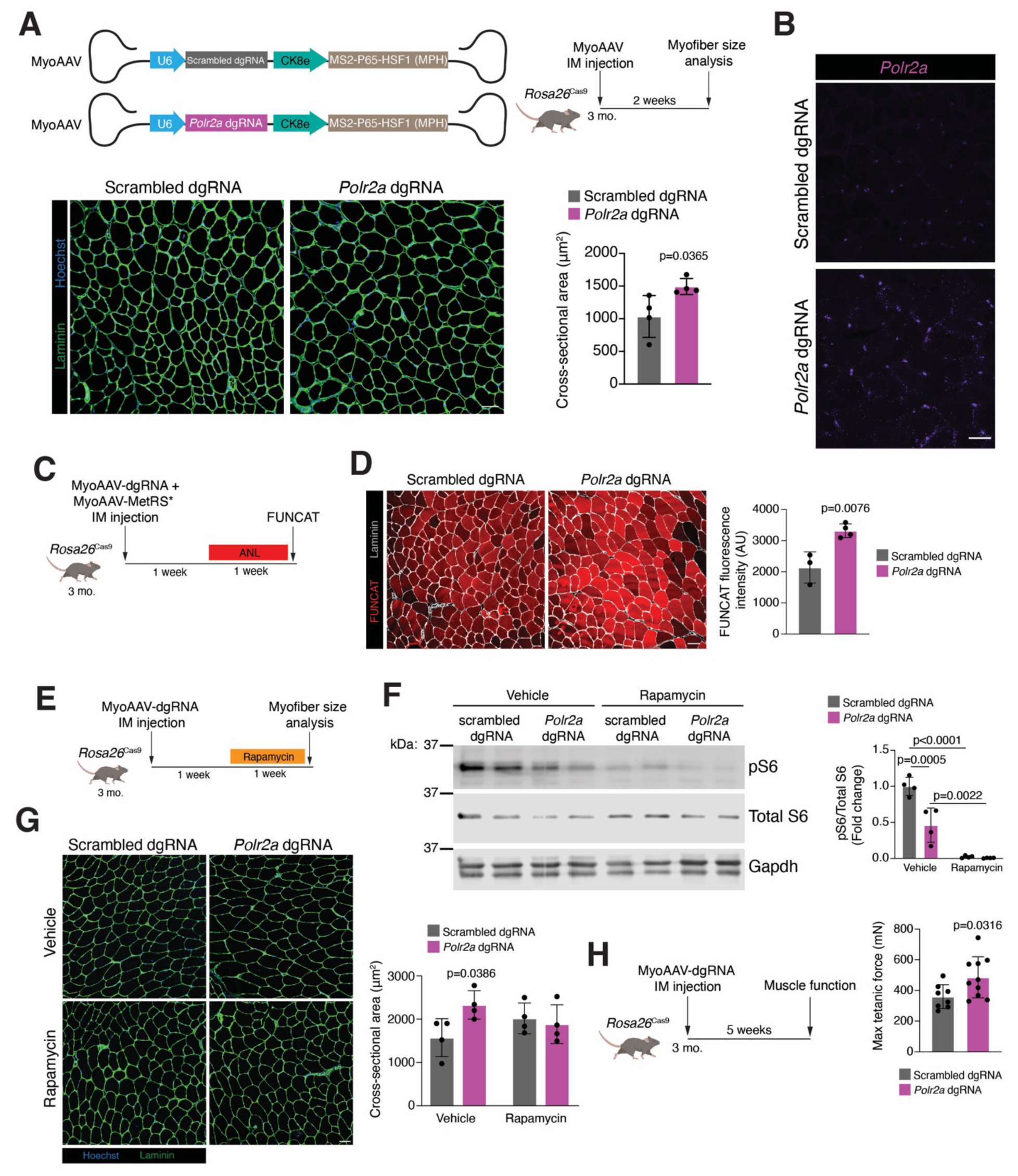
Regulation of functional muscle growth through activation of transcription in resident myonuclei. (A) Schematic representation of MyoAAV constructs containing scrambled or *Polr2a* dgRNA and the MPH complex under control of the CK8e promoter. *Rosa26*^Cas9^ tibialis anterior (TA) muscles were injected with the MyoAAVs. Representative images of TA sections immunostained with laminin antibodies and nuclei were labelled with Hoechst. Quantification of cross-sectional area of myofibers (n=4). Scale bar: 50 μm. (B) Representative images for *Polr2a* smRNA-FISH on TA muscles from adult *Rosa26*^Cas9^ mice. Scale bar: 20 μm. (C) Experimental setup for assessment of protein content using a mutated methionine-tRNA synthase (MetRS*) for fluorescent non-canonical amino acid tagging (FUNCAT). ANL is a non-canonical amino acid used by MetRS*. (D) Representative images of FUNCAT from TA sections after delivery of the indicated dgRNAs. Quantification of FUNCAT fluorescence intensity is shown on the right (n=3-4). Scale bar: 50 μm. (E) Schematic of experiment to analyze the effect of protein synthesis inhibition through rapamycin treatment on muscle growth after *Polr2a* upregulation. (F) Western blot for phospho-S6 ribosomal protein (pS6), total S6 protein, and Gapdh from TA lysates. Quantification of fold of change for the pS6/total S6 ratio is shown on the right (n=4). (G) Representative images of TA sections immunostained with laminin antibodies and nuclei were labelled with Hoechst. Quantification of myofiber cross-sectional area is shown on the right (n=4). Scale bar: 50 μm. (H) Experimental setup to assess muscle function. Quantification of max tetanic force generated is shown on the right (n=8-10). Data are presented as mean ± SD. Statistical test used were (A), (D) and (H) unpaired t-test; (F) two-way ANOVA with Tukey’s correction for multiple comparisons; (G) two-way ANOVA with Šídák’s correction for multiple comparisons.

We next characterized the quality of growth after *Polr2a* activation, specifically whether the increased myofiber volumes are associated with enhanced function. We determined whether myofibers with elevated mRNA transcription was associated with an increase in protein content by co-delivering a mutant methionyl-tRNA-synthetase (MetRS*) (*42–44*), which incorporates a non-canonical amino acid (ANL) into newly synthesized proteins, with the dgRNAs. We labelled newly synthesized proteins for 1 week after transduction with MyoAAV-MetRS* and either MyoAAV-CK8e-MPH-*Polr2a* dgRNA or MyoAAV-CK8e-MPH-scrambled dgRNA (Fig. 3C). Fluorescence intensity analysis in muscle sections revealed increased signal in muscles that received the *Polr2a* dgRNA (Fig. 3D), indicating that transcriptional increases are associated with accumulation of proteins. To test if protein synthesis is needed to drive growth after *Polr2a* upregulation, we treated mice with rapamycin (Fig. 3E), an inhibitor of mTOR-dependent protein synthesis (*45–47*). Rapamycin treatment resulted in reduced phosphorylated ribosomal S6 (pS6) subunit in both scrambled and *Polr2a* dgRNA muscle (Fig. 3F). We surprisingly detected reduced pS6 in *Polr2a* dgRNA muscle in the absence of rapamycin (Fig. 3F). Since we observed increased protein content in muscle with *Polr2a* overexpression (Fig. 3D), the reduced pS6 at baseline could suggest changes in the kinetics of protein synthesis after growth has been achieved. Consistent with increases in myofiber sizes being associated with elevated protein content, inhibition of protein synthesis through rapamycin treatment mitigated hypertrophy driven by *Polr2a* upregulation (Fig. 3G). The promotion of growth after *Polr2a* upregulation also led to elevated force production (Fig. 3H), overall indicating that an increase in transcription and mRNA content in myofibers is sufficient to promote functional muscle growth.

### Myonuclear transcriptional output uncouples accretion during muscle growth

Our data thus far indicate that *Polr2a* levels and mRNA transcription are limiting factors in the regulation of myofiber sizes and when activated in resident myonuclei can drive muscle growth. We next explored if one explanation for the need for the vast multinucleation in myofibers is related to mRNA content. If this paradigm is correct, one would also expect that elevated *Polr2a* in myofibers during times of robust myonuclear accretion may change the level of accretion. We thus upregulated *Polr2a* and enhanced transcription during two times of myonuclear accretion, postnatal developmental growth and adaptive growth in the adult in response to increased workload. MyoAAV-CK8e-MPH-*Polr2a* dgRNA or MyoAAV-CK8e-MPH-scrambled dgRNA was systemically delivered to *Rosa26*^Cas9^ mice at postnatal (P) day 6 (Fig. 4A) when fusion is ongoing in skeletal muscles (*2*). At P21 we observed a significant increase in *Polr2a* expression in tibialis anterior muscles (Fig. 4B). We also isolated single myofibers from EDL muscles and observed a reduction in the number of nuclei per myofiber (Fig. 4C). No differences were detected in EDL myofiber volume, and since myonuclear number was reduced, we found that myonuclear domain sizes (volume of cytoplasm/nucleus) are larger after *Polr2a* activation (Fig. 4D). We also observed that satellite cell numbers, as assessed by Pax7 immunostaining, are increased when *Polr2a* is upregulated in myofibers (Fig. S5A), consistent with fewer of these muscle progenitors being incorporated into the syncytium. In addition to postnatal development, myonuclear accretion is a hallmark of load-induced muscle growth in adult mice and humans (*6, 48–52*). To test if *Polr2a* upregulation impacts myonuclear accretion during adult hypertrophy, we utilized a genetic system we previously developed to track the newly fused nuclei (*41*). We crossed *Rosa26*^Cas9^ mice with *Pax7*^rtTA^; TRE^H2B-GFP^ mice, which allows labeling of satellite cell nuclei in a doxycycline-inducible manner. We then delivered MyoAAV-CK8e-MPH-*Polr2a* dgRNA or MyoAAV-CK8e-MPH-scrambled dgRNA intramuscularly into the lower hindlimb. After two days, we performed sham or tenotomy surgery to induce mechanical overload of plantaris muscles (*48, 53*). (Fig. 4E). Animals were treated with Doxycycline (Dox) during the entire duration of the study to activate H2B-GFP in satellite cells and allow tracking into myofibers. *Polr2a* was elevated in plantaris muscles that received MyoAAV-CK8e-MPH-*Polr2a* (Fig. 4F). While we observed an increase in myonuclear accretion in all the samples under tenotomy, as assessed by GFP^+^ myonuclei, muscles where *Polr2a* was elevated showed a reduced level of accretion (Fig. 4G). Despite reduced accretion after tenotomy in plantaris muscles where *Polr2a* is upregulated, an increase in cross-sectional area was observed (Fig. 4G). This data indicates that activation of transcriptional output of the resident myonuclei can support an increase in myofiber volumes of adult muscle while reducing the need for accrual of additional nuclei.

**Figure 4.**
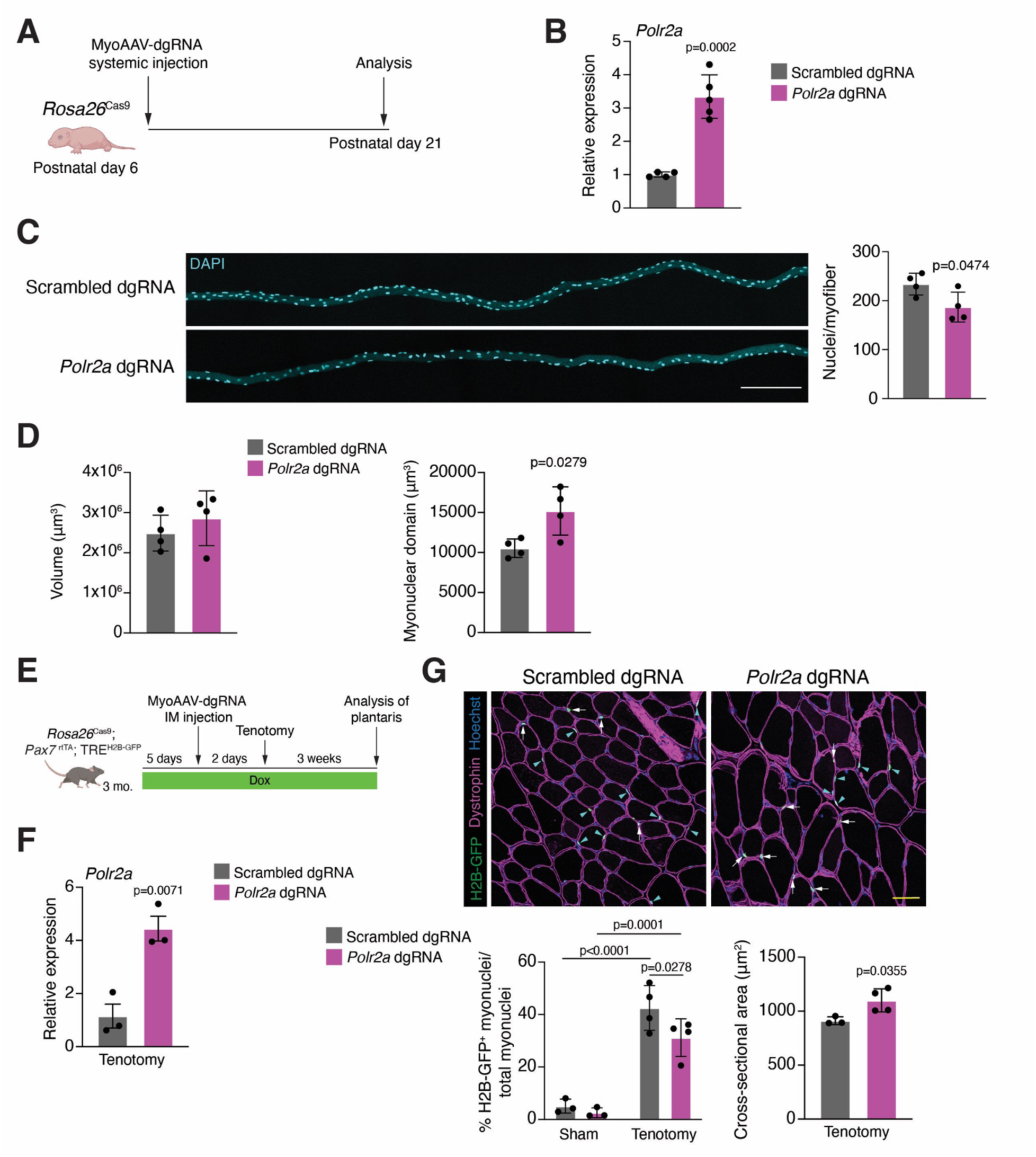
Increased transcriptional output in myofibers regulates accretion of activated satellite cells. (A) Experimental protocol followed to induce *Polr2a* upregulation in *Rosa26*^Cas9^ during postnatal development. Mice received MyoAAV systemically. (B) Quantitative RT-PCR for *Polr2a* expression in tibialis anterior muscles (n=4-5). (C) Representative images of segments of EDL myofibers isolated and stained with DAPI from *Rosa26*^Cas9^ mice treated with the indicated dgRNAs. Quantification of nuclei number per myofiber is shown on the right (n=4). Scale bar: 200 μm. (D) Quantification of volume (left) and myonuclear domain size (right) from myofibers shown in (C) (n=4). (E) Experimental protocol to assess nuclear accretion during adult muscle growth caused by tenotomy. *Rosa26*^Cas9^; *Pax7*^rtTA^; TRE^H2B-GFP^ mice were used to track accretion of myonuclei. (F) Quantitative RT-PCR for *Polr2a* expression in plantaris muscles under tenotomy from *Rosa26*^Cas9^; *Pax7*^rtTA^; TRE^H2B-GFP^ mice that received either scrambled or *Polr2a* dgRNA (n=3). (G) Representative images of plantaris muscles after tenotomy and immunostaining with dystrophin antibodies and nuclei were labelled with Hoechst. Arrows show H2B-GFP^+^ satellite cells and arrowheads highlight H2B-GFP^+^ newly fused myonuclei. Scale bar: 20 μm. Bottom, quantification of the percentage of H2B-GFP^+^ newly fused myonuclei normalized by the total number of myonuclei in plantaris muscles under sham and tenotomy conditions (n=3-4). Cross-sectional area of plantaris myofibers are shown after tenotomy (n=3-4). Data are presented as mean ± SD. Statistical tests used were (B), (D), (F) and (G, right graph) unpaired t-test; (G, left graph) two-way ANOVA with no correction for multiple comparisons.

## Discussion

Our results reveal that increasing transcriptional output in resident myonuclei is sufficient to cause myofibers to grow, even in the absence of any additional pro-growth stimuli. These findings advance our overall understanding of the links between mRNA transcription and cell size regulation. While mRNA content scaling with cell size during growth is established, and recent work has provided insight into how such scaling is regulated at the level of transcription (*21, 28*); however, the increase in myofiber size we report upon upregulation of Rpb1 shows that augmenting mRNA transcription and abundance within a cell can cause it to grow. This also indicates that RNAPII-mediated transcription is a limiting determinant of muscle growth. It is noteworthy that overexpression of just Rpb1, a single subunit of the RNAPII complex, results in increased mRNA transcription and cell size in myofibers. This contrasts with findings from yeast, where despite simultaneous elevation of multiple subunits of RNA polymerase II, there was no increase in cell size detected (*21, 28*). One reason for the difference in outcomes could be the duration of overexpression, which was acute in the yeast studies. Alternatively, terminally differentiated cells and more specifically multinucleated cells, such as myofibers, perhaps are more permissive for transcriptional induction of growth. This could be because they have more DNA content and transcriptional profiles dominated by transcripts encoding cytoskeletal and structural proteins whose expression directly impacts cell volumes. Indeed, we observed an increase in transcripts for several critical sarcomeric genes upon *Polr2a* overexpression, where the increase in size was associated with elevated total protein content and augmented force generation. Since our transcriptional analysis was restricted to specific candidate genes and did not take into account potential changes in transcriptome size, it is unclear if *Polr2a* overexpression globally altered the transcriptional profile within myofibers. Such a global increase in transcriptome size could diminish growth potential where resources are futilely utilized in the expression of genes and proteins not involved in growth.

A key discovery in our study is that the increase in myofiber size resulting from *Polr2a* activation does not involve an increase in myonuclear content of myofibers, meaning that no detectable fusion or nuclear accrual ensued when myofibers grew in response to increased transcription. This indicates that resident myonuclei within adult myofibers retain a reserve capacity to support larger myofiber volumes, which evidently becomes available for growth upon upregulation of Rpb1 expression. This was also found to be the case in a previously reported mouse model of restricted myonuclear accrual, where through titration of myonuclear numbers during postnatal development, we identified a transcriptional reserve capacity in resident myonuclei associated with increased levels of transcriptional activity (*17*). In *Drosophila*, resident myonuclei also exhibit reserve capacities, at the level of nuclei and nucleolar sizes, which can impact myofiber size and function (*54*). Increase in myofiber sizes without changes to myonuclear number has been reported in other experimental mouse models involving activation of Akt, JunB, or inactivation of myostatin (*55–57*). However, investigations in these models have focused on alterations in proteostasis, with increased protein synthesis and reduced protein degradation being identified as the principal driver of growth; whether increased myonuclear transcriptional output or other changes to the resident myonuclei also play a causal role in the myofiber growth reported in these models remains unknown.

This newly discovered link between increased RNAPII-mediated mRNA transcription within myofibers and myonuclear accrual was particularly evident under growth conditions that typically require fusion and accrual. Our findings that *Polr2a* upregulation reduces the number of myonuclei accrued during both postnatal developmental growth and adaptive growth in response to increased workload highlights a central role for myofibers in the regulation of satellite cell activity, fusion, and myonuclear accrual. More specifically, it suggests that the abundance or content of certain biosynthetic products within myofibers may regulate satellite cells and their fusion to myofibers. Global sarcomeric content, which in turn impacts myofiber size and function, might be one such influencing factor. Our data also raise the intriguing possibility that mRNA content within myofibers itself could be another factor dictating whether myofibers require, or accept, additional nuclei. If elevated mRNA content within myofibers is a negative regulator of myonuclear accretion, then mRNA dilution within myofibers could plausibly be a trigger for satellite cell activation and myonuclear accretion. Since satellite cells can be controlled through autonomous signaling mechanisms and through expression of factors on and coming from myofibers (*58–60*), one explanation could be that changes in myofiber mRNA content results in altered levels of niche factors or secreted factors that impact satellite cells. It is also possible that mRNAs in myofiber-derived extracellular vesicles can be sensed by satellite cells since there is evidence of satellite cell to myofiber communication through extracellular vesicles (*61*). Understanding any potential mRNA-dependent communication axis between myofibers and satellite cells will be essential to define how transcriptional scaling governs muscle homeostasis.

Growth of myofibers and maintenance of their volumes are known to be dependent on the number of nuclei and total protein abundance. This dependency on satellite cell accretion likely compromises the ability to maintain muscle size in pathological conditions such as sarcopenia, where satellite cells are impaired (*62–67*). We propose that enhancing the transcriptional capacity of existing myonuclei may bypass this limitation, providing a new layer of control over muscle size and function. It will be critical to evaluate the long-term consequences of sustained myonuclear transcriptional output, since aging can be associated with transcriptional alterations (*68–70*). Together, these results establish that the transcriptional capacity of resident myonuclei is a tractable process to control muscle mass and reveals a critical link between effects of increased myonuclear transcription and nuclear accretion in skeletal muscle, thus identifying an unanticipated paradigm that modulates skeletal muscle plasticity.

## Acknowledgements

We thank members of the Millay laboratory, Hima Bindu Durumutla for technical assistance and helpful discussions, and the Bio-Imaging and Analysis Facility at Cincinnati Children’s Hospital Medical Center.

## Author Contributions

D.P.M. acquired funding for the project. F.M.M., V.P., and D.P.M. conceived and designed the project. F.M.M., C.C., A.B., and A.A.W.C. performed experiments and analyzed data. C.O.S. analyzed the snATAC-seq data. X.L. and Y.D. built the snATAC-seq libraries. J.L.M. and R.J.K. performed and analyzed the muscle function experiments. All authors contributed to writing and editing the manuscript.

## Data and materials availability

All data are available in the manuscript or supplementary materials. Raw data and processed fragment files from snATAC-seq is accessible from NCBI Gene Expression Omnibus (GEO), GSE312647. Code for snATAC-seq analysis can be found at https://github.com/cswoboda/D2WATAC

## Materials and Methods

### Animals

To delete Myomaker in satellite cells and generate Δ2w mice, we interbred *Myomaker*^LacZ/+^ (*71*) (a null allele) with *Myomaker*^loxP/loxP^ mice (*72*) and also introduced a *Pax7*^CreER^ knock-in allele (*73*) that drives tamoxifen-inducible Cre recombinase specifically in satellite cells. Experimental mice were *Myomaker*^LacZ/loxP^; *Pax7*^CreER^ or *Myomaker*^loxP/loxP^; *Pax7*^CreER^, whereas littermates lacking Cre (*Myomaker*^LacZ/loxP^ or *Myomaker*^loxP/loxP^) served as controls. All mice received intraperitoneal injections of tamoxifen (Sigma-Aldrich) prepared in corn oil with 10% ethanol. Neonates were administered 200 µg of tamoxifen on postnatal day 6 to generate Δ2w mice as described previously (*17*). *Rosa26*^Cas9^ mice (*74*) were obtained from Jackson Laboratory (Strain #026179) and harbor ubiquitous Cas9 expression. *Pax7^rtTA^*; TRE^H2B-GFP^ mice were produced as previously described (*41*), and crossed to the *Rosa26*^Cas9^ line for myonuclear accretion studies. To induce H2B-GFP expression, animals were fed doxycycline (Dox) chow (0.0625%; TestDiet) according to the experimental timeline stated in the figures. All mice were maintained under standardized housing conditions (72°F, 30–75% humidity, 10/14 h light/dark cycle). All animal procedures were approved by the Institutional Animal Care and Use Committee at Cincinnati Children’s Hospital Medical Center.

### Muscle collection and preparation

Muscles were dissected and either snap-frozen in liquid nitrogen for molecular studies or embedded in tragacanth and frozen in liquid nitrogen–cooled 2-methylbutane for histology and stored at -80°C until further use. Samples allocated for histological assessment were cryosectioned at the mid-belly region and 10 μm sections were collected.

### snATAC-sequencing

The skeletal muscle snATAC-sequencing (snATAC-seq) libraries were prepared as described earlier (*75*). Briefly, frozen tibialis anterior muscles were cryosectioned on dry ice and homogenized using a gentleMACS Dissociator (Miltenyi; “Protein_01_01” program) in MACS buffer composed of 5 mM CaCl_2_, 2 mM EDTA, protease inhibitor (Roche), 3 mM MgAc, 10 mM Tris-HCl (pH 8.0), and 0.6 mM DTT. Homogenates were passed through a 30 μm CellTrics filter (Sysmex) and collected in 15 ml tubes. Nuclei were pelleted by centrifugation (500 × g, 5 min, 4°C), resuspended in 3 ml Nuclear Permeabilization Buffer (PBS with 5% BSA, 0.2% IGEPAL CA-630, 1 mM DTT, and protease inhibitors), and incubated on a rotating platform for 5 min at 4°C. After a second centrifugation, permeabilized nuclei were transferred into 500 μl of high-salt tagmentation buffer (36.3 mM Tris-acetate, 72.6 mM potassium acetate, 11 mM magnesium acetate, 17.6% DMF), counted by hemocytometer, and diluted to 1,000 nuclei in 9 μl.

Aliquots of 1,000 nuclei were distributed into four 96-well plates (384 wells total for 4 plates). Barcoded Tn5 transposomes (1 μl per well) were dispensed using a BenchSmart 96 (Mettler Toledo), and tagmentation proceeded for 90 min at 37°C with shaking (900 rpm). The reaction was stopped by adding 2.5 μl 100 mM EDTA (20 mM final) followed by an additional 15 min incubation at 37°C. Each well then received 6 μl of 3x sort buffer (PBS with 3% BSA and 3 mM EDTA). All wells were pooled, stained with Draq7 (1:500; Cell Signaling), and sorted on a Sony SH800 instrument. Eighty nuclei were deposited into each well of four new 96-well plates containing 8 μl of primer mix (15 pmol i7, 15 pmol i5, 200 ng BSA, and 0.027% SDS).

Next, plates were heated at 65°C for 25 min, after which 4 μl of 1.33% Triton X-100 was added to neutralize SDS. Each well then received 11 μl of Q5 PCR master mix (5 μl 5× buffer, 0.5 μl 10 mM dNTPs, 0.25 μl Q5 polymerase, and 5.25 μl water). Amplification was performed with the following cycling conditions: 72°C for 4 min; 98°C for 30 s; 12 cycles of 98°C for 10 s, 60°C for 30 s, 72°C for 45 s; and hold at 4°C. PCR products from all wells were pooled and purified using the Zymo DNA Clean & Concentrator-5 kit. Size selection of 200–600 bp fragments was performed on a 1.5% agarose gel, and DNA was recovered using the Zymo Gel DNA Recovery kit. Library concentration was measured using the Qubit 1x dsDNA HS Assay (Invitrogen), and fragment size distribution and nucleosomal periodicity were assessed on a High Sensitivity D1000 TapeStation (Agilent).

Next-generation sequencing of snATAC-seq libraries was performed on the DNBSEQ G400 platform using paired-end sequencing, 150-bp reads for each end, generating approximately 5,000–10,000 raw reads pair per cell. Raw read sequences were demultiplexed using cutadapt (https://cutadapt.readthedocs.io/en/stable/) and aligned to the 10-mm build of the mouse genome using snaptools (https://github.com/r3fang/SnapTools). Aligned bam files were converted to snap file format and then to sorted fragment files using snaptools’ snap-pre and dump-fragment functions, respectively.

snATAC-seq data were processed using the ArchR package (v1.0.3). Fragment files generated from snaptools were used to construct Arrow files with createArrowFiles(). Nuclei with less than 1000 minimum fragments, or a transcription start site (TSS) enrichment score of less than 4 were excluded from downstream analysis. Potential doublets were identified and excluded using addDoubletScores() followed by filterDoublets() with a filterRatio of 1.5. All arrow files, from all samples were then input into an ArchRProject using the ArchRProject() function. Dimensionality Reduction and Clustering Dimensionality reduction was performed using iterative Latent Semantic Indexing (LSI) via addIterativeLSI(), followed by clustering with the Seurat Leiden algorithm implemented in ArchR (addClusters()). Uniform Manifold Approximation and Projection (UMAP) embeddings were calculated with addUMAP(). Gene regulatory signals were quantified using ArchR’s gene score model using the addGeneScoreMatrix() function. Clusters were then labelled and merged into cell types based on gene score accessibility of known marker genes. These genes were visualized using the plotEmbedding() function, using imputed weights generated by the getImputeWeights() function which were provided to the imputeWeights parameter. These gene scores are shown in fig. S2.

In order to perform genome-wide peak-calling, cells were aggregated by cell type and condition using addGroupCoverages(). Genome-wide peak calling was carried out using MACS2 (v2.2.7.1) on pseudobulk replicates the pseudobulk replicates using the addReproduciblePeakSet() function, with the parameter .method = “Macs2”. MACS2 was executed with ArchR’s default ATAC-seq parameters to identify high-confidence, reproducible peaks across biological replicates. The resulting peak sets were merged to generate a unified peak-by-cell matrix via addPeakMatrix().

Differentially accessible peaks (DAPs) were identified between control and Δ2w myonuclei using getMarkerFeatures()with bias correction for TSS enrichment and fragment counts. Specifically, a binomial test was performed on a binarized peak matrix, using the parameters binarize = TRUE and testMethod = “binomial”. Once DAPs were generated, the list was filtered to retain only peaks with a log fold change of greater than or equal to 1, and an adjusted p-value of less than 0.01, generating a list of peaks more accessible in the Δ2w myonuclei. These peaks were then matched to the nearest gene annotated by MACS2 during peak calling. The nearest genes to these peaks (Data S1) were then input into ToppGene (https://toppgene.cchmc.org/). Selected gene ontology terms were then visualized based on number of genes present in that gene ontology term in a dot plot generated using the ggplot2 package.

### AAV production and delivery

The *Polr2a* dgRNA was designed in the portal CRISPick (broad.io/crispick) as a full-length gRNA (20 base pairs) for CRISPR activation. The full-length scrambled dgRNA was previously described (*76*). Based on these full-length gRNAs, we synthesized shortened versions (15 base pairs) to meet the criteria of dead gRNAs (*37, 77*) and the following sequences were used: *Polr2a* 5’-TTGACGCTAGGGCTC-3’, scrambled 5’-AAAGGAAGGAGTTGA-3’. Oligos were synthesized by IDT, annealed, and cloned into the pAAV-hU6-dgRNA-CAG-MPH plasmid (Addgene Plasmid #106259) (*37*). The hU6-dgRNAs and MPH genetic sequences were subcloned into a pAAV-CK8e plasmid for skeletal muscle-specific expression of MPH using In-Fusion cloning (Takara Bio). pAAV-GFP was obtained from Cell Biolabs, Inc. (Cat AAV-400).

AAVs were generated in-house using the MyoAAV-1A capsid (*38*), following established protocols. In brief, AAV Pro 293T cells (Takara) were seeded in 15 cm plates at 2x10⁷ cells per dish and transfected the following day with 10.5 μg of helper plasmid, 5.25 μg of the MyoAAV-1A Rep/Cap plasmid, and 5.25 μg of the expression construct. Recombinant virus was collected from both cells and supernatant and purified by iodixanol-gradient ultracentrifugation (*78*). Viral genome titers were determined by qPCR using a C1000 Touch Thermal Cycler (Bio-Rad). For *in vivo* delivery, AAVs were administered by intramuscular injection into hindlimb muscles at 1x10^11^ vg, or by intrathoracic injections at 5x10^10^ vg.

### Immunofluorescence and cross-sectional area (CSA) analysis

Muscle cryosections were fixed in 4% PFA/PBS for 10 min, washed three times with PBS, permeabilized with 0.5% Triton X-100/PBS for 10 min, washed three times with PBS, and blocked with 3% BSA/0.025% Tween-20/PBS for 1 h at room temperature in a humidity chamber. Slides were then incubated with rabbit anti-laminin (1:300; Sigma cat. #L9393) or rabbit anti-dystrophin (1:100; Abcam cat. #ab15277) primary antibodies for 3 h at room temperature, or overnight at 4°C, in a humidity chamber. For Pax7 staining, antigen retrieval was performed as previously described (*79*) on 10 μm TA muscle cryosections. After antigen retrieval, cryosections were blocked with 3% BSA/0.025% Tween-20/PBS for 90 min followed by a second blocking with 3% BSA/0.025% Tween-20/PBS supplemented with Mouse-On-Mouse IgG Blocking Reagent (Novus Biologicals cat. #MKB-2213-NB) for 90 min at room temperature. Then sections were incubated with anti-Pax7 antibodies (1:10; Pax7, Developmental Studies Hybridoma Bank) overnight at 4 °C, in a humidity chamber. After incubation with primary antibodies, slides were washed three times with PBS and incubated with goat anti-rabbit (1:500; Invitrogen) and goat anti-mouse IgG1 (1:300; Invitrogen) Alexa Fluor secondary antibodies in blocking buffer for 90 min at room temperature in a humidity chamber. Slides were then washed three times with PBS and one time with water. 1 μg/mL of Hoechst 33342 (Sigma-Aldrich) was added to counterstain nuclei. Fluoromount-G (Southern Biotech cat. #0100-01) was used for mounting. Representative images were taken on a Nikon A1R confocal microscope for use in CSA analysis. The CSA of individual myofibers were measured using NIS Elements software (Nikon), which can quantify the area within laminin-labeled myofibers.

### Myofiber isolation and nuclear counting

EDL muscles were isolated and digested in high-glucose DMEM (HyClone) containing 0.2% collagenase type I (Sigma-Aldrich) at 37°C. After 40–45 min, tissues were gently triturated with a wide-bore glass pipette to release individual myofibers. Isolated myofibers were transferred to high-glucose DMEM supplemented with 10% horse serum and maintained at room temperature. Myofibers were then washed three times in PBS and fixed in 4% PFA for 10 min at room temperature, followed by three additional PBS washes and storage in PBS at 4°C. For nuclear staining, myofibers were mounted onto Superfrost Plus slides (Thermo Fisher Scientific) using VectaShield with DAPI (Vector Laboratories) and coverslipped. Images were acquired on a Nikon A1R confocal microscope. Myonuclear number and myofiber length were quantified in Imaris Bitplane 10.2 (Oxford Instruments). Myofiber volume was calculated by averaging diameters measured at five positions along each myofiber, determining cross-sectional area, and multiplying by length.

### smRNA-FISH on cryosections and EDL myofibers

Single-molecule RNA-FISH was carried out using the RNAscope platform (ACDBio) according to the manufacturer’s guidelines on 10 μm fresh-frozen muscle cross-sections. In brief, cryosections were fixed with 4% PFA/PBS at 4°C for 15 min followed by a dehydration step with an ethanol (EtOH) series of 50% EtOH for 5 min, 70% EtOH for 5 min and 100% EtOH twice for 5 min at room temperature. After that, cryosections were stored in 100% EtOH at -20°C for not longer than a week. Right before perform smRNA-FISH, the muscle cryosections were air drayed and treated with hydrogen peroxide for 15 min at room temperature, washed twice with PBS, and followed by protease IV treatment for 30 min at room temperature in a humidity control tray, followed by two washes with PBS. Probes were hybridized at 40°C for 2 h, and amplification and HRP development steps were performed according to RNAscope kit instructions. For smRNA-FISH on isolated EDL myofibers, we followed the protocol described by Kann and Krauss (*80*). Myofibers were isolated, fixed in 4% PFA/PBS for 10 min at room temperature, washed three times in PBS, transferred to 100% methanol, and stored at −20°C until further use. Prior to staining, myofibers were rehydrated through a graded methanol-to-PBST (PBS + 0.1% Tween-20) series (50% MeOH/50% PBST; 30% MeOH/70% PBST; 100% PBST) for 5 min each on a rotator at room temperature. Rehydrated myofibers were mounted on Superfrost Plus slides (Thermo Fisher Scientific) and treated with Protease III for 15 min at room temperature, followed by three washes with PBST. Probes were hybridized overnight at 40°C, and amplification and HRP development steps were performed according to RNAscope kit instructions. For nuclear staining, cryosections and myofibers were mounted using VectaShield with DAPI (Vector Laboratories) and coverslipped.

Representative z-stacks images were taken on a Nikon A1R confocal microscope and signal intensity was quantified in Fiji by measuring the mean gray value within defined regions of myofibers or tissue sections. Transcript numbers were determined using rendered images in Imaris Bitplane 10.2 (Oxford Instruments) by counting discrete mRNA spots within defined fiber segments.

### mRNA levels in fibroblast syncytia

To assess total mRNA levels in multinucleated cells, 10T½ fibroblasts were transduced with lentiviruses containing a GFP construct and a Myomaker and Myomerger doxycycline-inducible expression cassette. These cells were seeded in 8-chamber Ibidi slides at 4x10^4^ cells/chamber on day 0 and incubated in DMEM containing 10% Tet-Free fetal bovine serum (Peak Serum, cat. #PS-FB2) overnight. On the next day, Myomaker and Myomerger expression was induced by treating cells with doxycycline (Dox) containing complete medium (1 μg/mL), and fusion was evaluated 24 h after Dox treatment. Next, cells were fixed with 4% PFA and smRNA-FISH was performed using polyA and GFP probe using the RNAscope platform (ACDBio) according to the manufacturer’s guidelines. Representative z-stacks images were taken on a Nikon A1R confocal microscope. Images were then converted for use in Imaris Bitplane 10.2 software (Oxford Instruments). Cell surfaces were generated by selecting a region of interest using the GFP channel as a proxy for cell volume. Channel intensity was measured per 3D surface cell volume.

### Tenotomy

Tenotomy surgery was performed as previously described (*48*). In brief, an incision on the lower limb’s posterior-lateral side was made to expose the soleus and gastrocnemius muscles. The distal and proximal tendons of the soleus were cut followed by cutting of the distal gastrocnemius tendon and removal of 1mm of the tendon to avoid regeneration. After the surgery, the mouse leg was sutured and maintained according to the experimental timeline shown in the figure.

### Protein synthesis and protein content assessment

Protein synthesis was quantified using fluorescent noncanonical amino acid tagging (FUNCAT) as previously described (*81*). MetRS*, a mutant methionyl-tRNA synthase, was subcloned from a pMaRSC construct (Addgene plasmid # 89189) (*82*) into pAAV. *Rosa26*^Cas9^ mice were transduced with 1x10^11^ vg of a MyoAAV-MetRS* through intramuscular injection of the tibialis anterior muscles. One week later, mice were provided drinking water containing 10 mM ANL H-L-Lys(N₃)-OH·HCl (Iris Biotech HAA1625). ANL was dissolved in autoclaved tap water supplemented with 0.7% maltose and filtered through a 0.22 μm membrane and provided to mice *ad libitum* for 7 days as described (*44*). Tibialis anterior muscles were collected and FUNCAT was performed on 10 μm muscle cryosections. In brief, cryosections were fixed with 4% PFA/PBS for 15 min, washed twice with 3% BSA/PBS, permeabilized with 0.5% Triton X-100/PBS for 20 min and washed twice again with 3% BSA/PBS at room temperature. The Click chemistry reactions was performed using the Click-it EdU Imaging Kit (C10646, ThermoFisher Scientific) and replacing the Alexa Fluor picolyl azide dye with TAMRA-alkyne (900932, Sigma-Aldrich) for 30 min at room temperature and protected from light. After that, the reaction cocktail was removed and cryosections washed twice with 3% BSA/PBS followed by immunostaining with laminin antibodies as describe above. Representative images were taken on a Nikon A1R confocal microscope and FUNCAT intensity was measured using NIS Elements software (Nikon).

### Rapamycin administration

Rapamycin (LKT labs, cat. #53123-88-9) was dissolved in DMSO to make a 5 μg/μl stock solution. For *in vivo* experiments, mice received 2 mg/kg rapamycin in 100 μl of PBS via intraperitoneal injection every 24 h for 7 days. For vehicle conditions, mice received an equivalent amount of DMSO dissolved in 100 μl of PBS.

### Max tetanic force analysis

Muscle performance was measured *in situ* with a 300C-LRmuscle lever system (Aurora Scientific Inc., Aurora, CAN). Briefly, the mice were anesthetized via inhalation (∼3% isoflurane, SomnoSuite, Kent Scientific) and placed on a thermostatically controlled table with anesthesia maintained via nose-cone (∼2% isoflurane). The knee was firmly fixed and the skin overlying the lower hindlimb partially dissected. The tibialis anterior tendon was isolated and tied with a silk suture t0 the motor shaft. Contraction of the tibialis anterior was elicited by percutaneous electrical stimulation of the peroneal nerve (0.2 msec pulse duration) and optimal length was determined using single twitches spaced apart to prevent fatigue. A force-frequency response was performed from 1 hz to 150 hz and maximal tetanic force at 150 hz determined.

### Immunoblot analysis

Snap-frozen muscles were resuspended in 800 μl RIPA lysis buffer (50 mM Tris-HCl with pH 7.5, 150 mM NaCl, 2 mM EDTA, 1 mM MgCl_2_, 0.1% SDS, 1 mM dithiothreitol, 1 mM sodium orthovanadate, 1 mM Benzamidine, 1% NP-40 and protease inhibitors) and minced with scissors for 3 min until homogenize. Samples were then digested for 1 h in rotation at 4°C and then lysates were sonicated on low intensity three times for five seconds each. Muscle lysates were centrifugate on a table-top centrifuge at 12,000 rpm for 10 min at 4°C. The supernatants were transferred to a new tube and protein concentration was measured by BCA assay (ThermoFisher) and 50 μg was heated in 4X Laemmli buffer at 95°C for 5 min. Tissue lysates were fractionated by 10% SDS-PAGE gels, transferred onto a PVDF membrane (ThermoFisher cat. #88518), stained with Ponceau S (Thermo Scientific cat. #A40000279) for total protein visualization and immunoblotted using the following antibodies: Rpb1 CTD (Cell signaling technology, cat. #2629; 1:500), Phospho-S6 Ribosomal Protein (Ser240/244) (Cell signaling technology, cat. #5364; 1:1000), S6 Ribosomal Protein (5G10) (Cell signaling technology, cat. #2217; 1 ;1000), Gapdh (Millipore, cat. #MAB374; 1;5000) in 1% BSA in TBS buffer with 0.05% Tween-20. Secondary antibodies conjugated to AlexaFluor-680 (Invitrogen, cat. #A21102), DyLight 680, or DyLight 800 (Cell Signaling Technology, cat. #5470S or #5151P) were used for detection. Protein bands were visualized using the Odyssey imaging system (LI-COR Biosciences) and analyzed for densitometry using Fiji.

### Gene expression analysis

Total RNA was isolated from muscles samples using TRIzol^TM^ reagent (ThermoFisher) according to the manufacturer instructions. 0.5-2 μg of total RNA was used to synthesize cDNA using the Superscript III Kit (Invitrogen 18080400) with oligo dT primers. Primers were designed using the NCBI primer-blast tool and commercially synthesized (Custom DNA Oligos; IDT). Semi-quantitative RT-PCR was carried out on Biorad CFX 96 using the PowerUp SYBR Green Master Mix (Applied Byosistems), with a reaction volume of 20 μL, primer concentration of 0.8 μM and cDNA concentration of 20 ng. Data normalization was done by the 2^−ΔΔCt^ method using the average of the reference gene as described previously (*83*). Primer sequences: *Polr2a* forward, 5’-CTGGACCTACCGGCATGTTC-3’; reverse, 5’-GTCATCCCGCTCCCAACAC-3’, *Myh4* forward, 5’- CTTTGCTTACGTCAGTCAAGGT-3’; reverse, 5’-AGCGCCTGTGAGCTT GTAAA-3’, *Acta1* forward, 5’-CTACTGAGCCACACGCCAG-3’; reverse, 5’- TCACACACAAGAGCGGTGG-3’, and *Gapdh* forward, 5’-TGCGACTTCAACAGCAACTC-3’; reverse, 5’-CCTCTCTTGCTCAGTGTCC-3’

### Total and nascent RNA transcription analysis

Total and nascent transcription was measured by RNA labeling using the Click-iT Nascent RNA capture kit (Invitrogen, cat. # C10365) according to the manufacturer’s instructions. In short, control and Δ2w mice were treated with 1 mg of 5-ethynyluridine (EU) through intraperitoneal injections at postnatal day 59. After 6 h of EU pulse, tibialis anterior muscles were harvested, and total RNA was extracted using the TRIzol^TM^ reagent (ThermoFisher). This RNA was then used in a copper-catalyzed click reaction with an azide–biotin to biotinylate RNA containing EU (nascent transcripts). The biotin transcripts were captured on streptavidin magnetic beads and the non-biotin labelled transcripts were used as total RNA. These RNA pools were used for cDNA synthesis using the Superscript III Kit (Invitrogen 18080400) with oligo dT primers. Quantitative RT-PCR was performed as described above and relative transcript levels were determined from threshold cycles for amplification using the 2^−ΔΔCt^ method and normalized by the myonuclear number average per myofiber from control or Δ2w mice.

### Statistical analysis

Sample sizes are noted in the figure legends. For isolated myofibers, at least 10 myofibers per animal were analyzed and data are plotted as means per animal, except for fig. S4B where individual myofibers from three independent animals is plotted. Data were compared between groups using various statistical tests, which are indicated in the figure legends, along with specific p-values. First, the data were assessed for normality using a Shapiro-Wilk test. For comparison of two groups, an unpaired t-test was used unless an F test indicated significantly different variances between the two groups, in which case an unpaired t-test with Welch’s correction was used. A Mann-Whitney test was used for comparison of two groups when a group did not pass the Shapiro-Wilk test. For comparison of three or more groups, with one independent variable, we used a one-way ANOVA with Tukey’s correction for multiple comparisons. For comparison of three or more groups, with two independent variables, we used a two-way ANOVA with Tukey’s or Šídák’s correction for multiple comparisons. For data in Fig. 4G (left graph) specific comparisons were determined a priori, thus we used a two-way ANOVA with no correction for multiple comparisons. In all graphs, error bars indicate standard deviation (SD). Data were processed using GraphPad Prism 9 software.

---

**Figure S1.**
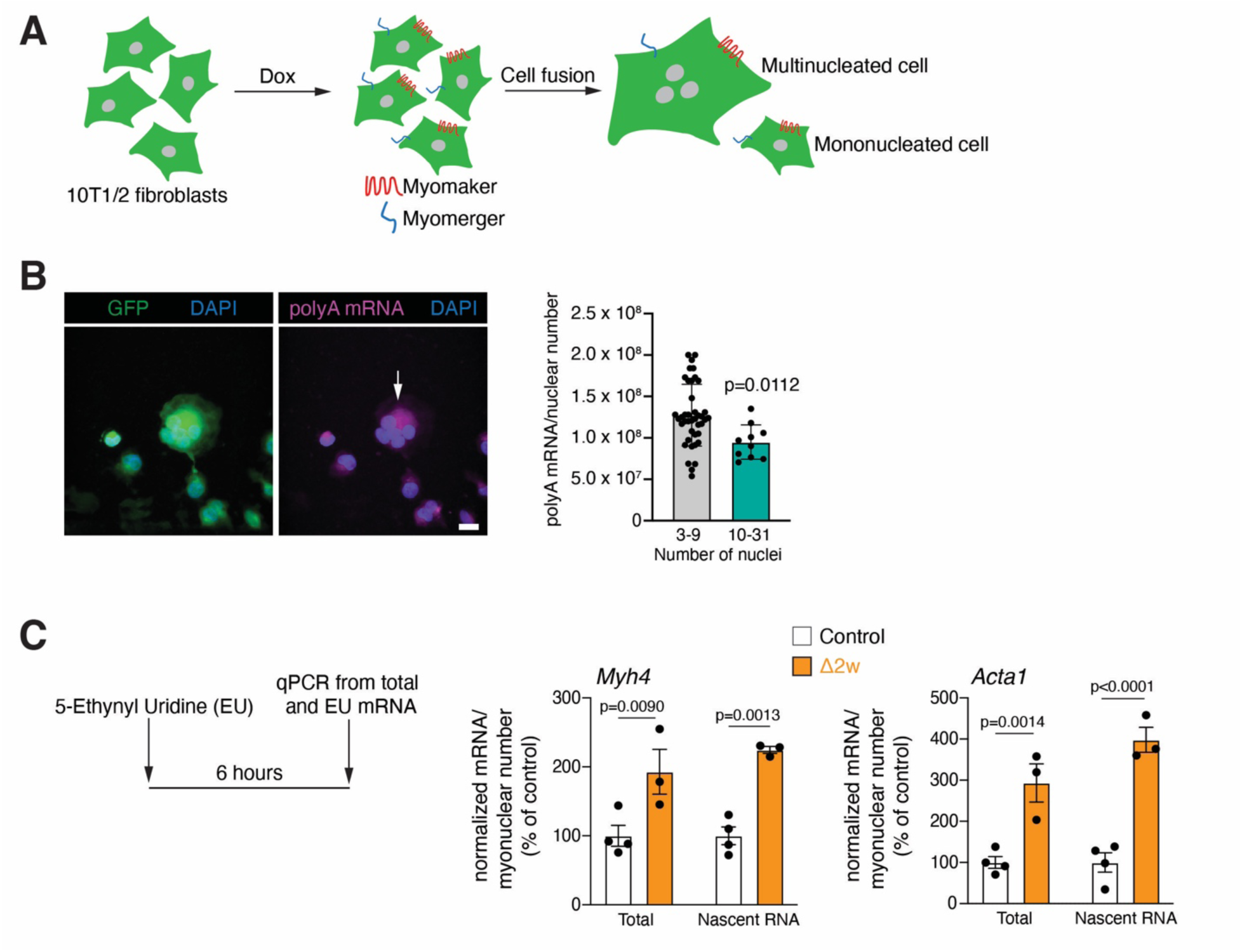
mRNA levels and transcription in syncytial cells. (A) Experimental setup to generate artificial syncytiums through Dox-inducible expression of Myomaker and Myomerger in fibroblasts. (B) Representative smRNA-FISH images for polyA mRNA in multinucleated 10T ½ fibroblasts (arrow). The fibroblasts were genetically labeled with GFP. Scale bar: 10 μm. Quantification of polyA mRNA intensity normalized by nuclear number in fibroblasts with 3-9 or 10-31 nuclei (n=individual cells). (C) Experimental protocol to analyze active transcription in control and Δ2w mice. Quantitative RT-PCR for *Myh4* and *Acta1* from total and nascent RNA (EU-labelled RNA) (n=3-4). Data are presented as mean ± SD. Statistical tests used were (B) unpaired t-test; (C) two-way ANOVA with Šídák’s correction for multiple comparisons.

**Figure S2.**
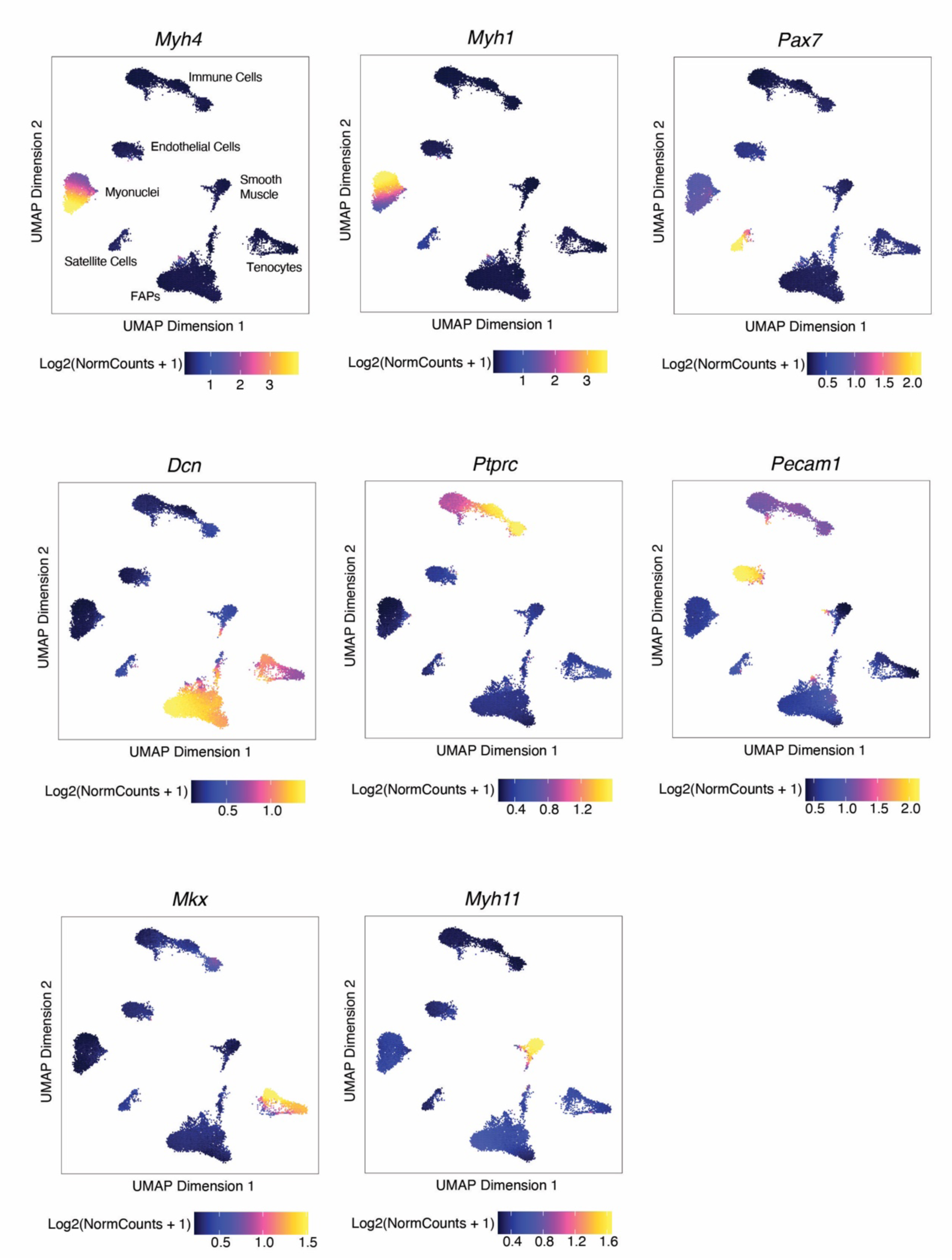
UMAP representation of gene accessibility scores for known marker genes of cell populations. From snATAC-seq data, gene accessibility scores for *Myh4* (myonuclei), *Myh1* (myonuclei), *Pax7* (satellite cells), *Dcn* (FAPs), *Ptprc* (immune cells), *Pecam1* (endothelial cells), *Mkx* (tenocytes), and *Myh11* (smooth muscle) were used to identify cell populations. Cell population identities are shown in the *Myh4* UMAP.

**Figure S3.**
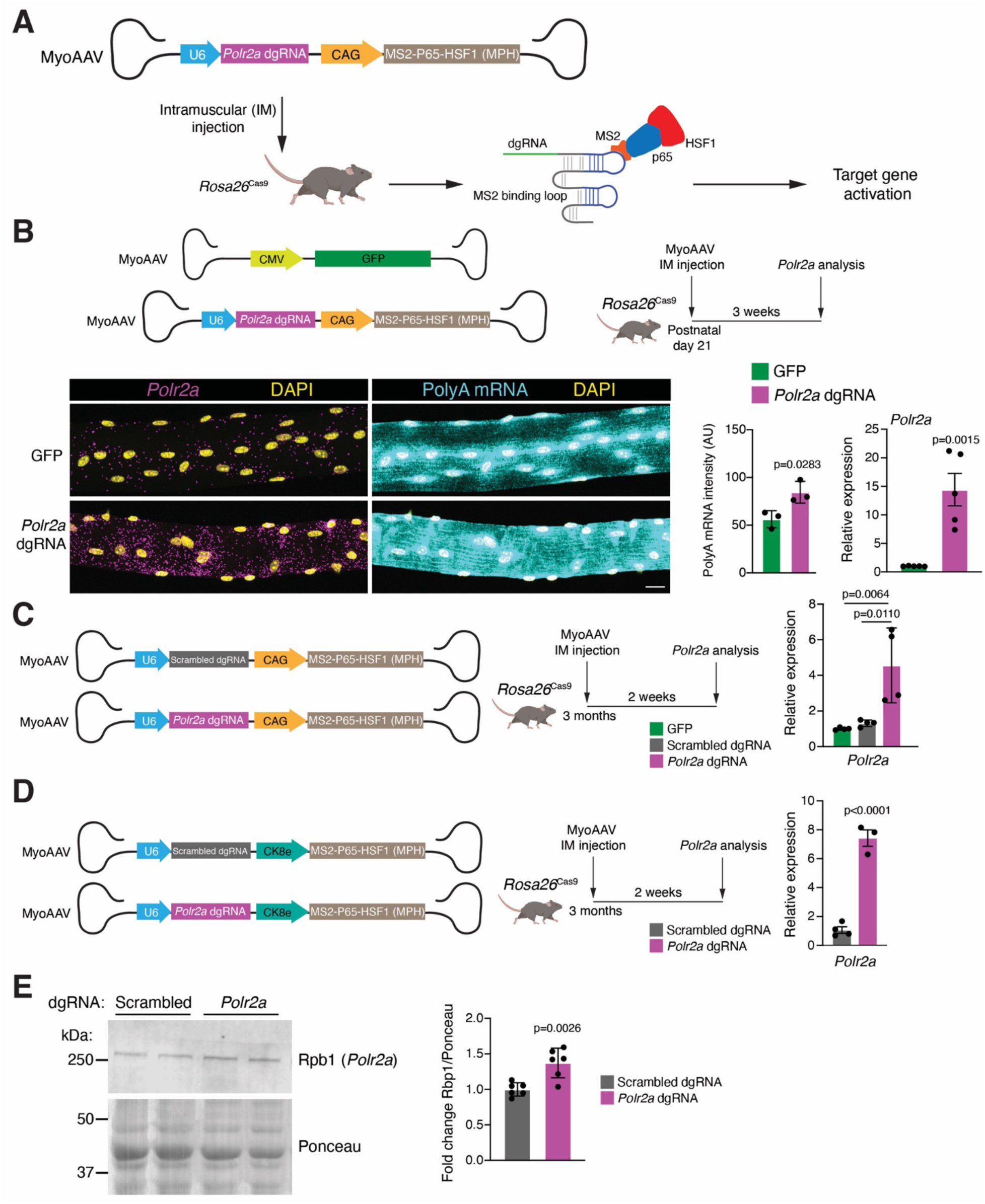
Multiple models can induce *Polr2a* upregulation and transcriptional output in myofibers *in vivo.* (A) Schematic representation of the target gene activation system used to upregulate *Polr2a* expression in *Rosa26*^Cas9^ mice. (B) Schematic of the MyoAAV constructs and protocol used to transduce *Rosa26*^Cas9^ tibialis anterior (TA) muscles with MyoAAV-CMV-GFP or MyoAAV-CAG-MPH-*Polr2a* dgRNA. Representative smRNA-FISH images for *Polr2a* and polyA mRNA from EDL myofibers. Scale bar: 10 μm. Quantification of polyA mRNA intensity (n=3) and quantitative RT-PCR for *Polr2a* expression in tibialis anterior muscles from *Rosa26*^Cas9^ mice transduced with MyoAAV-CMV-GFP or MyoAAV-CAG-MPH-*Polr2a* dgRNA (n=5). (C) Schematic of MyoAAV constructs and experimental protocol. Quantitative RT-PCR for *Polr2a* expression in tibialis anterior muscles transduced with the indicated MyoAAVs (n=4). (D) Schematic of MyoAAV constructs and experimental protocol. Quantitative RT-PCR for *Polr2a* expression in tibialis anterior muscles transduced with MyoAAV-CK8e-MPH-scrambled dgRNA or MyoAAV-CK8e-MPH-*Polr2a* dgRNA (n=3-4). (E) Western blot for Rpb1 (product of *Polr2a* mRNA) and Ponceau staining as a loading control from tibialis anterior lysates from mice used in (D). Quantification of fold of change for Rpb1/Ponceau ratio (n=6). Data are presented as mean ± SD. Statistical tests used were (B), (D) and (E) unpaired t-test; (C) one-way ANOVA with Tukey’s multiple comparisons test.

**Figure S4.**
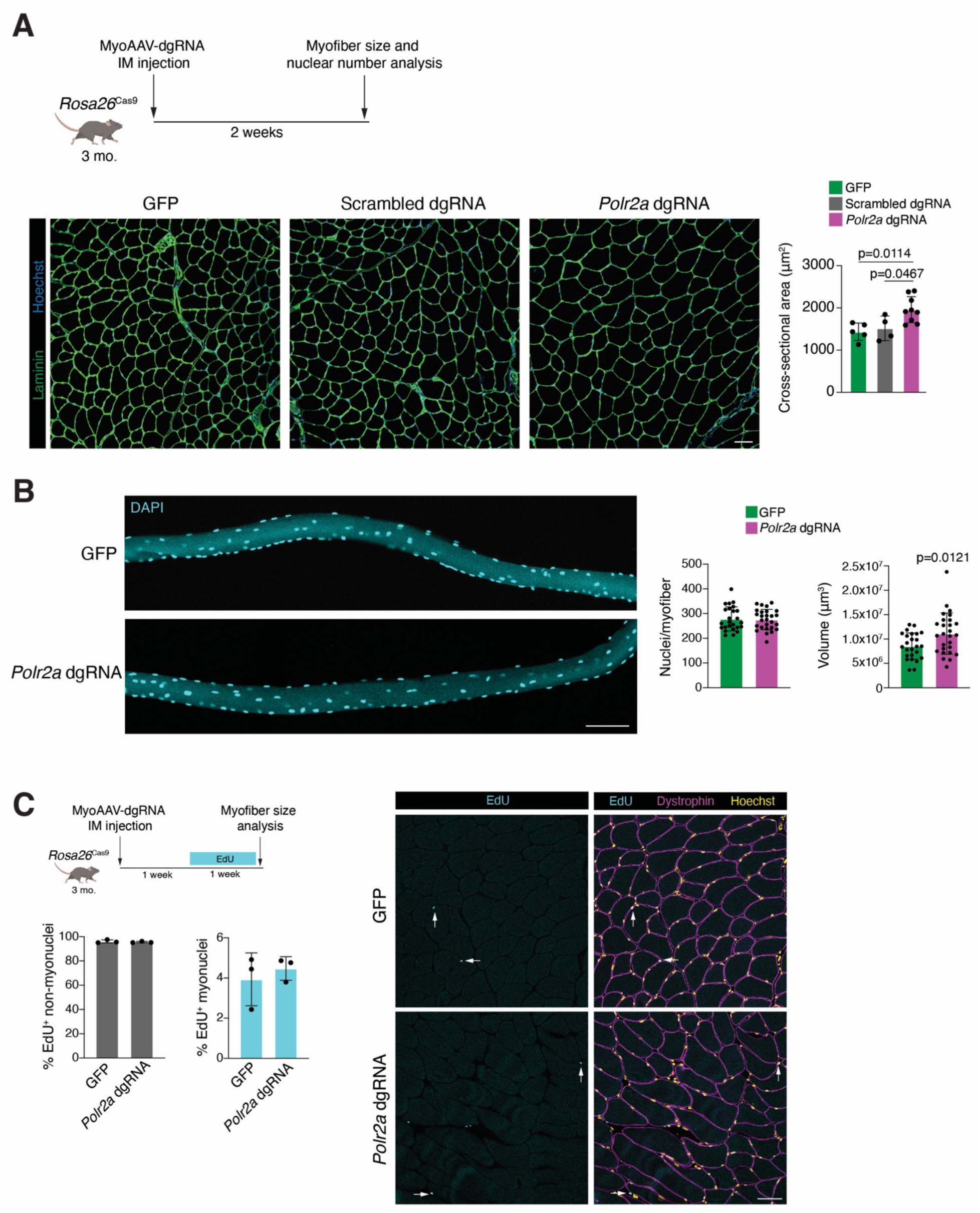
Upregulation of *Polr2a* and transcriptional output in myofibers increases muscle size independent of myonuclear accretion. (A) Experimental protocol followed to upregulate *Polr2a* expression in myofibers. Representative images of tibialis anterior sections immunostained with laminin antibodies from *Rosa26*^Cas9^ mice transduced with MyoAAV-CMV-GFP, MyoAAV-CAG-MPH-scrambled dgRNA or MyoAAV-CAG-MPH-*Polr2a* dgRNA. Nuclei are labelled with Hoechst. Quantification of cross-sectional area is also shown (n=4-9). Scale bar: 50 μm. (B) Representative images of EDL myofibers isolated from 3-month-old *Rosa26*^Cas9^ mice transduced with MyoAAV-CMV-GFP or MyoAAV-CAG-MPH-*Polr2a* dgRNA. Quantification of nuclei number per myofiber and volume from the indicated myofibers (n=25-27 myofibers from 3 independent animals). Scale bar: 100 μm. (C) Schematic representation of protocol used to track satellite cell proliferation and accretion into myofibers. 3-month-old *Rosa26*^Cas9^ mice were transduced with MyoAAV-CMV-GFP or MyoAAV-CAG-MPH-*Polr2a* dgRNA and treated with EdU for 7 days prior to sacrifice. Representative images of tibialis anterior sections immunostained with dystrophin antibodies. Nuclei were labelled with Hoechst and EdU^+^ nuclei through Click-It chemistry. Arrows show EdU^+^ nuclei. Scale bar: 50 μm. Quantification of the percentage of EdU^+^ non-myonuclei and EdU^+^ myonuclei is shown on the left (n=3). Data are presented as mean ± SD. Statistical tests used were (A) one-way ANOVA with Tukey’s correction for multiple comparisons; (B) unpaired t-test with Welch’s correction.

**Figure S5.**
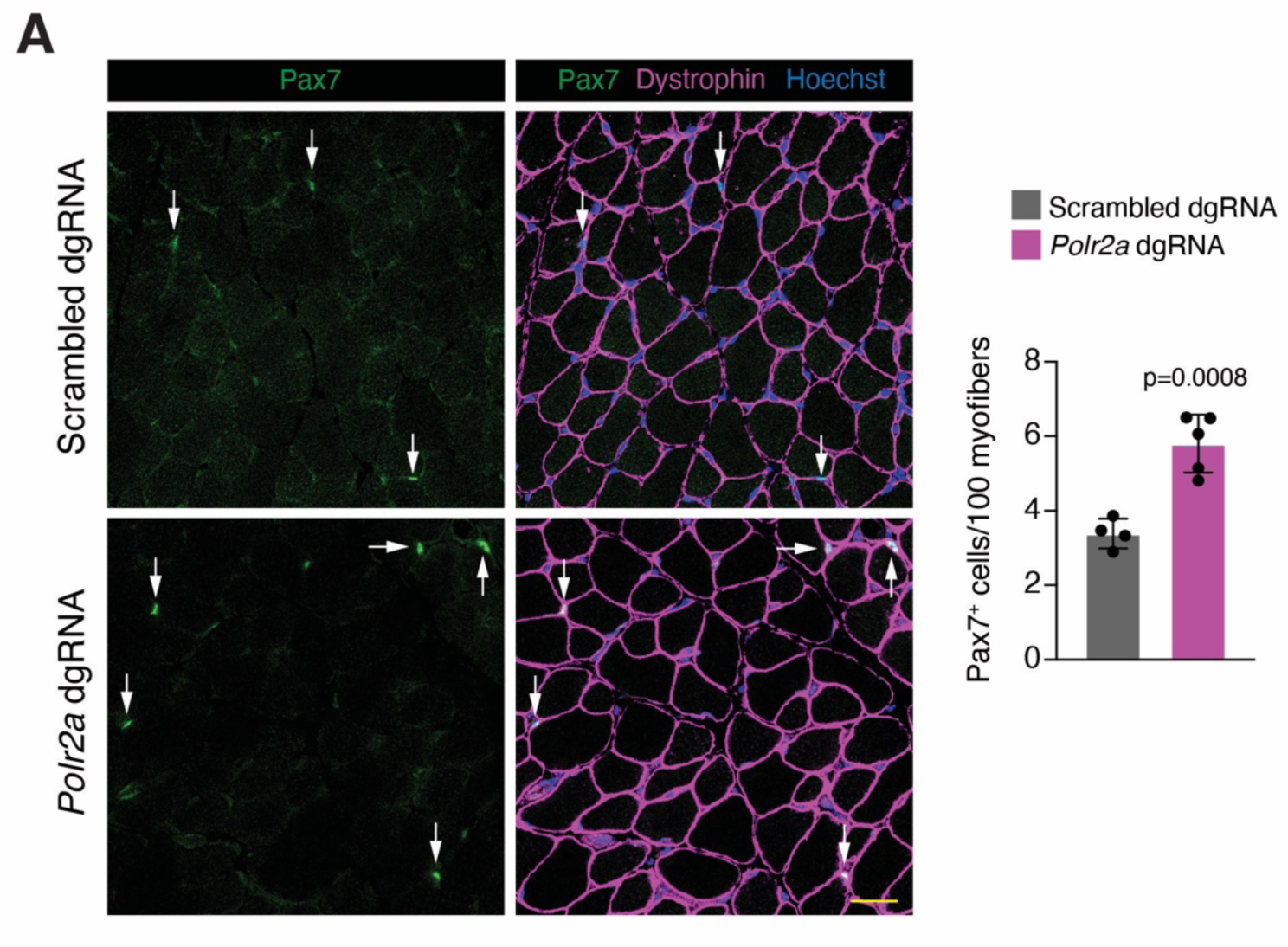
Satellite cell numbers are increased after myofiber-specific *Polr2a* upregulation during postnatal development. (A) Representative images of tibialis anterior sections from *Rosa26*^Cas9^ mice transduced with MyoAAV-CK8e-MPH-scrambled dgRNA or MyoAAV-CK8e-MPH-*Polr2a* dgRNA at postnatal day 6 and analyzed at postnatal day 21. Sections were immunostained with Pax7 and dystrophin antibodies and nuclei were labelled with Hoechst. Arrows show Pax7^+^ cells. Scale bar: 10 μm. Quantification of Pax7^+^ cells per 100 myofibers is shown on the right (n=4-5). Data are presented as mean ± SD. Statistical test used was unpaired t-test.

